# High-throughput microCT scanning of small specimens: preparation, packing, parameters and post-processing

**DOI:** 10.1101/2020.01.22.911875

**Authors:** Christy A. Hipsley, Rocio Aguilar, Jay R. Black, Scott A. Hocknull

## Abstract

High-resolution X-ray microcomputed tomography, or microCT (μCT), enables the digital imaging of whole objects in three dimensions. The power of μCT to visualise internal features without disarticulation makes it particularly valuable for the study of museum collections, which house millions of physical specimens documenting the spatio-temporal patterns of life. Despite its potential for comparative analyses, most μCT studies include limited numbers of museum specimens, due to the challenges of digitising numerous individuals within a project scope. Here we describe a method for high-throughput μCT scanning of hundreds of small (< 2 cm) specimens in a single container, followed by individual labelling and archival storage. We also explore the effects of various packing materials and multiple specimens per capsule to minimize sample movement that can degrade image quality, and hence μCT investment. We demonstrate this protocol on vertebrate fossils from Queensland Museum, Australia, as part of an effort to track community responses to climate change over evolutionary time. This system can be easily modified for other types of wet and dry material amenable to X-ray attenuation, including geological, botanical and zoological samples, providing greater access to large-scale phenotypic data and adding value to global collections.

## Introduction

High-resolution X-ray microcomputed tomography, also known as HRXMT or microCT (μCT), is an increasingly powerful tool for the non-destructive investigation of whole objects. Functioning like a microscope with X-ray vision, μCT generates high fidelity 3D models of solid material from which the outer layers can be virtually dissected or removed, revealing the inner structure. Starting with radiographs of an object taken over multiple angles, a computer algorithm is used to digitally reconstruct a stack of 2D X-ray projections, or tomograms, into a 3D volume. Whereas in human medicine the X-ray source rotates around the patient (e.g., computerized axial tomography, or CAT scan), in μCT the object is typically fixed on a rotating stage while the X-ray tube remains stationary. The differential properties of the object’s matter, including thickness and atomic number, interact with the X-ray’s energy beam to determine the number of photons that pass through it to reach the detector on the other side. This decrease in electromagnetic radiation, termed X-ray attenuation, results in detector pixels with grayscale values proportional to the radiopacity of the material, meaning that dense regions such as bone or rock appear white or light gray (radiopaque), while muscle or skin appears dark (radiolucent). The improved resolution of μCT over standard imaging techniques can achieve a detail detectability down to 200 nm (0.2 micrometre) – less than the diameter of a single red blood cell.

Since publication of the first X-ray microtomographic figures nearly four decades ago^1,2,3^, μCT has had profound impacts across scientific disciplines. Studies in biomedicine, zoology, geology and paleontology now regularly incorporate μCT images^4,5,6^, and open source software for the quantitative analysis of volumetric data is developing rapidly (e.g., Fiji^7^, Blob3D^8^, Dragonfly [Object Research Systems (ORS) Inc., Montreal, Canada]). Within the biological sciences, μCT has been particularly valuable for the study of museum collections, which contain millions of often small, delicate and unique specimens not amenable to traditional (destructive and/or irreversible) preparation. Worldwide, these collections span taxonomic, geographic and temporal distributions, providing a wealth of information for understanding the past, present and future of biodiversity^9^. Creation of cybertypes, or virtual models of type material, is another emerging application of μCT technology^10,11^, allowing researchers around the world to interact with voucher specimens. When combined with downstream analyses like geometric morphometrics, these models can be used to test taxonomic hypotheses that could shorten the lagtime between species discovery and formal description^10,12^, thus expanding the taxonomic bottleneck.

Despite the increasing use of μCT in systematic biology^13^, a major challenge remains in the practical imaging of high numbers of museum specimens within a project scope, for example in the context of large-scale analyses of phenotypic variation. This has proven difficult because of the need for samples to remain motionless during scan time (minutes to hours), and because each individual must be digitally labelled to match the physical specimen’s identity and hence retain important information, e.g., material type, locality, stratigraphic age. These issues have so far limited community-level μCT analyses, which could provide important insights into evolutionary responses to environmental change in small, rare, and/or fossilized taxa^6^. While other mass digitisation efforts of museum collections are already under way, they are typically limited to high resolution photographs of entire drawers to capture external morphology^14,15^. However internal structures are among the fastest responders to environmental drivers, suggesting an untapped role for μCT data in climate change research^6^. Especially in the Anthropocene, μCT offers a promising digital technique to identify the evolutionary consequences of human activities in natural history collections at both the internal organismal and ecosystem levels^16,17^.

Although some digital morphology studies have included hundreds of^18,19^ or over a thousand^20,21,22,23^ 3D models, their images are often aggregated using different sources (museums, repositories, laboratories), methods (surface scanning, μCT), and parameters (X-ray energy, voxel size, number of projections), making it difficult to integrate and compare findings. For smaller specimens in particular, variation in spatial resolution can result in significant measurement errors, for example in estimated volumes or relationships between traits^24,25^. Limitations on time and money for μCT scanning also mean that specimen data are often collected haphazardly, constraining systematic analyses to one or few individuals per species^6^. This trade-off between data accessibility and data quantity (and quality^26^) precludes the opportunity for more rigorous investigations of phenotypic diversity in time and space, as well as focused analyses of variation in a single taxon or locality. Common biodiversity metrics such as species identity, richness, and evenness are inferable from μCT data, e.g.,^27,28^, especially for cryptic taxa and species complexes with few observable differences^29,23,31^. Therefore, optimisation of the scanning process for high resolution 3D images of multiple specimens would not only contribute to our understanding of how biodiversity responds to global change, but also to the value of museum collections and researchers’ abilities to access them.

Here we outline steps for the high-throughput μCT scanning of small (< 2 cm) fossils, meant to facilitate advanced exploration of museum collections and allow researchers with limited access to specialized facilities the opportunity to maximize their investments. In contrast to other high-throughput μCT methods that have focused on clinical evaluations^32,33^ or automated phenotyping of laboratory strains (e.g., mice^34^, rice^35^, zebrafish^36^), we concentrate on the practical arrangement of accessioned museum specimens so that each individual can be identified and labelled in the 3D volume, followed by long-term storage and archiving. We illustrate this method using vertebrate (reptile, amphibian and mammal) fossils from the paleontological collections of Queensland Museum, Australia, as part of a larger effort to identify changes in morphology and species assemblages over geological time. We succeed in maximizing the quantity of fossils per container to over 200 specimens to generate high quality 3D models for comparative analyses, while minimising associated time, handling, and error. These steps should be applicable to any small dry objects amendable to X-ray attenuation including geological material, plant tissues, and invertebrates, and wet specimens provided they can be mounted inside small capsules or other containers^37,38^.

## Methods

### Preparation

The first step is the only time the user is required to handle specimen material directly, we therefore recommend the use of featherweight spring or feather light forceps and secondary containment (see Figure 1 for an overview of packing material). Although specimens of varying size can be scanned together, it is preferable to organise fossils into batches of roughly similar material properties for X-ray optimization. Thin, unbuffered, neutral pH tissue is ideal for specimen packing and eventual long-term storage. Other required materials are small clear two-piece pharmaceutical capsules, paper and/or plastic drinking straws, archival paper and pen for labelling, a 50 ml plastic centrifuge tube with cap, and medium density polyethylene foam. Suggested suppliers and estimated costs of material per tube are listed in Table 1.

**Figure 1.**
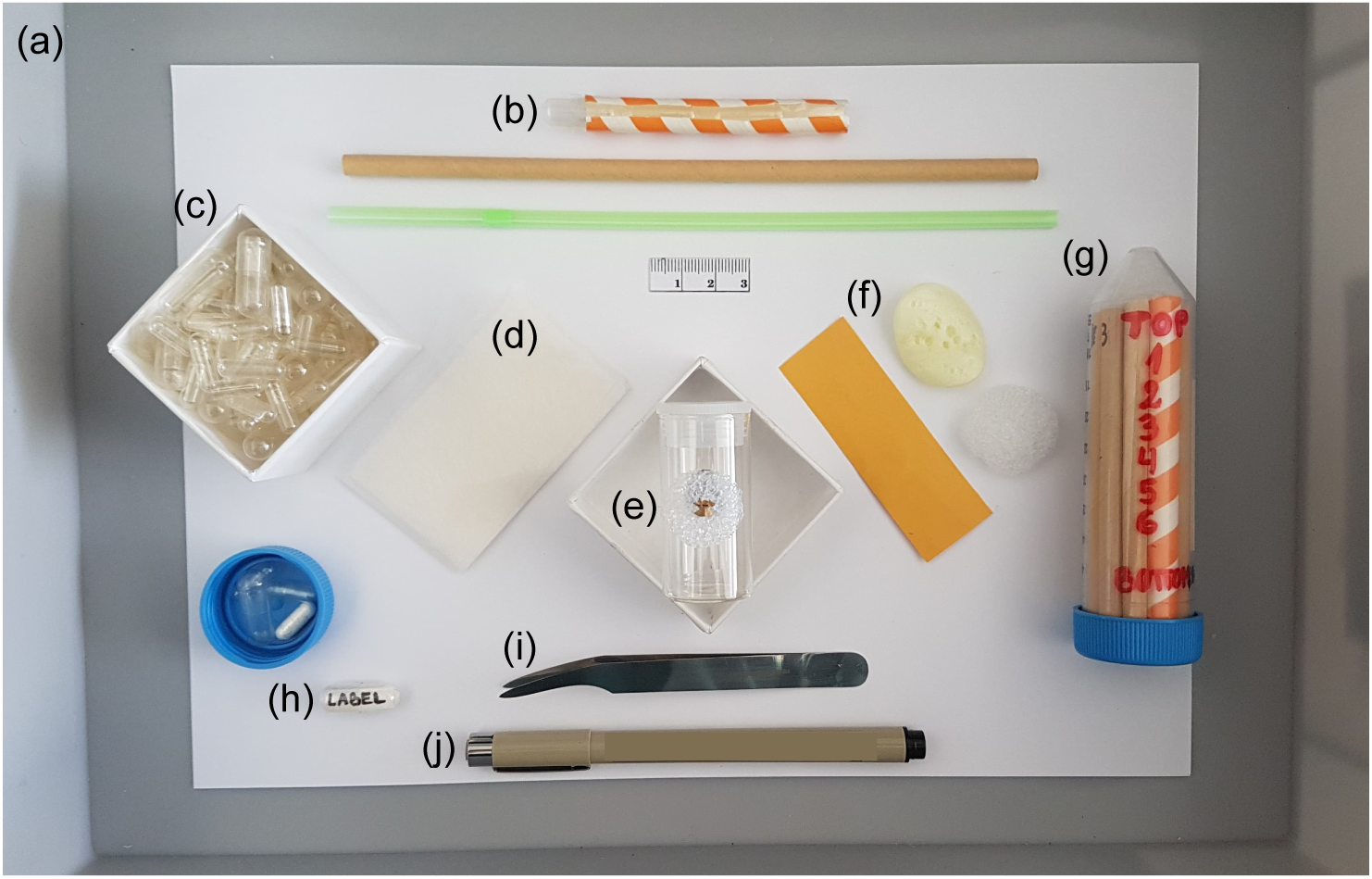
Equipment for dense fossil packing prior to μCT scanning: (a) litter tray, (b) paper and/or plastic straws, (c) clear two-piece pharmaceutical capsules, (d) archival tissue, (e) dry fossil specimen, (f) supporting material (paper, foam), (g) centrifuge tube with cap, (h) archival paper for labels shown inside a size 4 capsule, (i) forceps, (j) archival pen. Estimated costs and material suppliers are listed in Table 1.

**Table 1.** Estimated costs in Australian dollars (AUD) for material to μCT scan a 50 ml cylinder fully packed with 84 capsules holding 1 small (< 1 cm long) specimen each. Note that many items (e.g., Falcon tube, straws, forceps, foam, litter tray, pen) can be reused multiple times, thus decreasing the cost for subsequent scans.

| Item | Total cost per scan (AUD) |
| --- | --- |
| Centrifuge Falcon Tube 50 mL polypropylene (PP), conical bottom with screw cap<sup>1</sup> | 0.60 |
| Hard 2-piece clear gelatine capsules, size 4 (volume 0.23 ml)<sup>2*</sup> | 2.10 |
| Standard paper straw (6 x 135 mm)<sup>3</sup> | 0.27 |
| Green's Lens Tissue, un-buffered TIS-L Soft, acid-free, long fibred 9gsm repair/wrapping tissue (609 x 914 mm)<sup>4</sup> | 0.76 |
| Feather Forcep Sharp E122S, 0.23 mm gauge thick<sup>5</sup> | 4.90 |
| Ethafoam 220, medium density, non-cross linked Polyethylene (PE) closed cell foam (2400 x1200 mm)<sup>6</sup> | 1.00 |
| Litter tray<sup>7</sup> | 4.00 |
| Label paper<sup>8</sup> | 0.10 |
| Archival pen<sup>9</sup> | 6.54 |
| TOTAL | 20.30 |

Suppliers: <sup>1</sup>https://corning.com.au; <sup>2</sup>https://gelcapsules.com.au; <sup>3</sup>https://www.greenpack.com.au;
<sup>4</sup>http://www.archivalsurvival.com.au; <sup>5</sup>http://www.entosupplies.com.au;
<sup>6</sup>https://www.foamsales.com.au; <sup>7</sup>https://www.kmart.com.au; <sup>8</sup>https://www.talasonline.com;
<sup>9</sup>https://www.newtownartsupplies.com.au/
\*Clear 2-piece vegetable capsules are also available, although we did not test their performance in μCT scans.

### Packing procedure

Briefly, each specimen is wrapped in archival tissue and placed inside a capsule with its label, several of which are inserted lengthwise into a straw. Up to 14 standard paper straws cut to 9 cm in length fit inside a closed 50 ml Falcon tube, with each straw holding 6 size 4 capsules. For easier tracking of specimens in the 3D volume, we recommend leaving one capsule empty in different positions along the length of some straws (see post-processing section below). Following this method, over 80 specimens < 1 cm long can be scanned in a single tube. This quantity can be doubled or tripled when 2 or 3 fossils (separated by tissue) are packed inside the same capsule, allowing a maximum of 252 specimens. However, scans with more than 3 specimens per capsule showed a high (≥ 50%) rate of movement during rotation, compared to those with fewer (Table 2). We therefore limit packing to 1-2 specimens per capsule, provided they can be distinguished from one another in relative size and/or shape. Specimens up to 2 cm long can be similarly packed inside size 00 capsules. Larger diameter paper or smoothie straws sliced lengthwise hold 3 size 00 capsules in a 50 ml tube (Fig. 1b), which can be scanned together with smaller straws as needed.

**Table 2.** Estimated movement among 408 fossils μCT scanned using various packing materials to secure the specimen inside each capsule.

| Packing material | Number of specimens per capsule (size 4) that moved during scanning |  |  | Archival? | Comments |
| --- | --- | --- | --- | --- | --- |
|  | 1-2 | 3 | > 3 |  |  |
| Green's Lens<sup>1</sup> | 0/100<br>(0%) | 2/39<br>(5.1%) | 25/50<br>(50%) | Yes | Ideal consistency |
| Kim Tech wipes<sup>2</sup> | 0/100<br>(0%) | 4/39<br>(10.3%) | 27/50<br>(54%) | No | 100% fibre, not acid free |
| Ethafoam 220<sup>3</sup> | 10/10<br>(100%) | 10/10<br>(100%) | 10/10<br>(100%) | No | Slippery and hard |
| Tyvek<sup>4</sup> | - | - | - | Yes | Not tested, too waxy |
| Renaissance<sup>5</sup> | - | - | - | Yes | Not tested, too rigid |

Suppliers: <sup>1</sup>http://www.archivalsurvival.com.au; <sup>2</sup>https://www.keprofessional.com.au;
<sup>3</sup>https://www.foamsales.com.au; <sup>4</sup>http://www.dupont.com.au; <sup>5</sup>http://www.lightimpressionsdirect.com

Labels for each specimen are written in Indian or archival ink on uncoated acid-free paper, providing a unique and durable identifier that will be matched in the 3D volume. Our labels followed a two-part system containing specimen ID (e.g., museum accession number) and a code for straw and capsule position. Using permanent marker, each straw is labelled with a letter (A, B, C…N for 14 straws) and each capsule is labelled with its straw’s letter and a number from 1-6, denoting its position in the straw from top to bottom (Fig. 2a). This system can be amended to suit user needs, for example by including abbreviations for taxon if known and/or material (e.g., ‘il’ for ilium, ‘max’ for maxilla). These codes can also be extracted from the digital labels and used as variables in downstream analyses, e.g., in the R statistical package geomorph^39^ or MorphoJ^40^. Record labels with capsule and straw positions, including any empty capsules as a marker, since these will be checked off later during unpacking.

**Figure 2.**
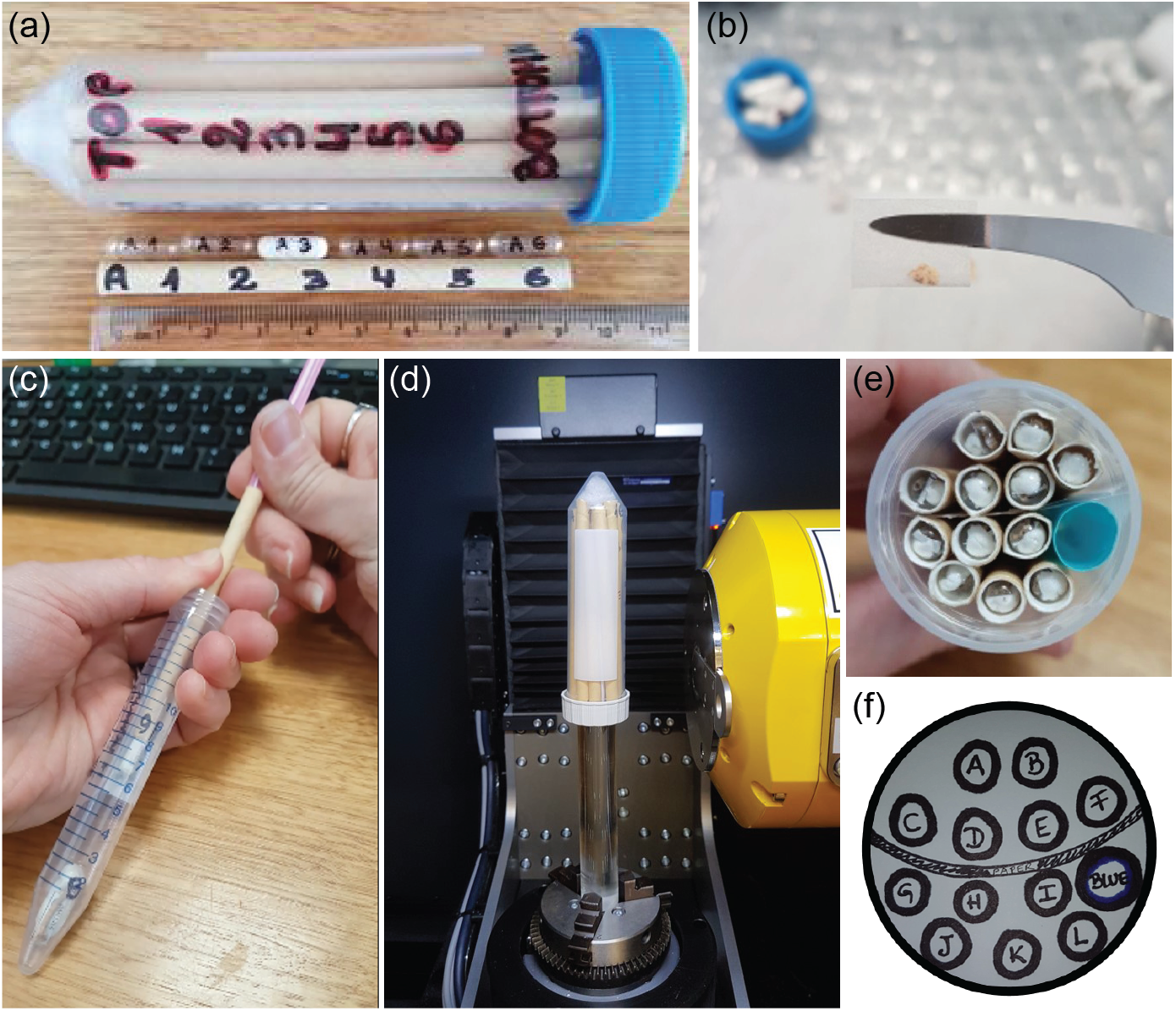
Specimen labelling and packing: (a) 50 ml Falcon tube with a labelled straw and capsules, (b) a single fossil wrapped in Green’s Lens tissue to minimise movement during image acquisition, (c) loading capsules into the paper straw using a thinner cocktail straw to gently push them down, (d) a packed 50 ml Falcon tube mounted on a glass rod inside the μCT machine; note that the cap-side is facing down (e) the packed tube from above, and (f) the diagram of straw arrangements in (e), noting the position of a larger blue straw and paper used as a divider.

Once labelling is complete, the specimens, labels, capsules and straws are aligned for packing. Each specimen is wrapped in a small envelope of Green’s Lens tissue (Fig. 2b), and empty spaces in the capsule are filled with tissue so the specimen does not contact its walls. This creates more distance between adjacent samples, allowing easier separation (i.e. segmentation) in the 3D volume. The tissue also prevents movement during image acquisition (Table 2), which can lead to motion artefacts in the reconstruction like shadows or streaking. Finally, the label is inserted into the capsule with the ID facing out for long-term storage, and the capsules are loaded into their respective straws in the correct order. A thinner cocktail straw can be used to gently push the capsules down without compressing ones below it (Fig. 2c).

To load the packed straws into the Falcon tube, first fill the conical tip with polyethylene or other firm material to create a flat surface. Insert the straws with position 1 towards this end, which will be facing up in the μCT machine. The flat surface of the cap on the other end can be fixed to a glass rod or dowel using a hot glue gun to elevate the tube from the machine’s stage (Fig. 2d). This avoids including the metal clamp in the scan which could affect X-ray optimization, and centres the tube vertically and horizontally relative to the detector. Draw a diagram of the straw arrangement with the list of specimen labels, noting where paper or empty straws are used to separate them for easier identification (Fig. 2e,f). This material will be visible in reconstructed cross-sections, allowing one to match the 3D volume to the diagram. Finally, medium density foam can be cut and placed as filler around the straws and under the cap of the tube before closing. This foam is recommended over tissue or other supporting material as it acts as an effective shock absorber during rotation. Secure the tube onto the stage, making sure it can rotate freely without touching the X-ray source or detector.

### X-ray parameters

Scan settings vary depending on the equipment, material size and properties, and desired results. Here we report parameters as optimised on a Phoenix nanotom m (GE Sensing & Inspection Technologies GmbH, Wunstorf, Germany) equipped with a 180 kV, 20 W high-power nanofocus X-ray tube and DXR detector (3072 × 2400 pixels). As with all microtomographic imaging, initial calibration of the detector is largely software-driven to reduce potential noise due to defective pixels or incident light. We calibrated the detector by taking an offset image (dark field) and two sets of gain images (flat field) at the same energy of the scan and at half the current, where images were averaged over 100 snapshots of the detector with a skip of 10. For all scans our instrument was fitted with a tungsten target, although molybdenum may be preferable for amber^4^. Fossils can be radiodense, in such cases reconstruction artefacts may occur when lower energy X-rays are attenuated at the surface, or when stronger X-rays hit the detector without passing through the object first (beam hardening)^4^. A thin metal filter placed in front of the X-ray source can overcome these issues by removing lower energy photons, although other parameters like current, voltage, or exposure time should be modified to achieve appropriate contrast.

Ideally the number of images for reconstruction is equal to the maximum width of the scanned object in pixels × π/2^38,41^. For a sample optimized to fill the full width of our detector, this would result in nearly 5,000 projections at a resolution of 10.6 μm. Such a data set would be massive in terms of scan time and storage (up to 70 GB before reconstruction), in addition to the computing power and time needed to reconstruct and render each of the 3D files individually. To improve the μCT process for smaller specimens without sacrificing image quality, we divided the Falcon tube into three segments by moving the position of the stage up or down between scans, while keeping all other settings identical. This produced separate volumes of equally high quality while avoiding the need to move the detector back and/or the tube forward to accommodate its length, thereby decreasing resolution. These volumes can be merged using Avizo (Thermo Fisher Scientific) or other dedicated software to recreate the entire tube in high resolution. However, for faster processing we aligned each segment to include two rows of size 4 capsules (or one row of size 00), so they could be separately visualised and labelled in the 3D volume if preferred.

For all reconstructions we followed the standard protocol of the Phoenix datos|x software (GE Sensing & Inspection Technologies GmbH, Wunstorf, Germany), which tests for object movement by comparing the first and last 2D projections. At high resolution, even minute movements can cause visible artefacts in the reconstructed images^4,38^. Instead of correcting for movement of single specimens during reconstruction by shifting the axis of rotation (which would in turn generate artefacts in the rest of the tube), we identified and labelled those specimens in post-processing so they could be rescanned. Other settings applied during reconstruction were a 3×3 median filter and an ROI filter.

### Computed tomography post-processing

Following reconstruction, digital labelling of the specimens is performed in 3D volume rendering software. Here we describe steps using VGStudioMax 3.1 (Volume Graphics, Heidelberg, Germany), although other options for free or commercial licenses are available (see Table 7 in Keklikoglou et al.^38^). To match the tube orientation to the diagram (Fig. 2f), the volume can be rotated in the 3D window or by using the registration tool in 2D. Straws can then be labelled digitally with the indicator tool or by creating a region of interest (ROI) to name each one by its letter (A, B, C… N). Similar-sized capsules should be aligned across the straws horizontally, making it easy to identify capsule positions in the 2D and 3D windows. For example, starting from the top (tip) of the tube in straw A, specimens should be ordered as A1, A2…A6, with capsules in the neighbouring straw B following the same order. Verifying the positions of empty capsules is another safeguard to ensure correct orientation. Using ROIs, the specimens can be segmented and renamed in the rendering software to match the specimen labels (e.g., museum accession number plus straw/capsule position). False-colouring straws or capsule rows differently in the 3D volume also helps to track specimen order. Any specimens with obvious artefacts can be marked with their labels followed by the word ‘REDO’, to be set aside during unpacking so they can be scanned again under improved conditions, i.e, repositioned in the capsule or with better tissue support.

### Unpacking and storage

Capsules are unpacked from the straws in the same manner as they were put in, by inserting a cocktail straw into one end and pushing it through (Fig. 2c). We recommend checking specimens off the list as they are being unpacked as a final test of correct labelling. These samples can be stored long-term inside the capsules since archival tissue and paper was used. Specimens for rescanning can be included in the next batch and labelled digitally with their ID, new straw and capsule position, and the word ‘REDONE’ for transparency. This process can be repeated as many times as necessary to achieve optimal quality scans for all material. Likewise, all labelled straws and Falcon tubes can be reused in subsequent scans, which will further ensure consistency in the packing and labelling process.

## Results and Discussion

Optimal settings for our fossils in terms of time, resolution and file size were three separate scans of the Falcon tube at 40 mm focus-to-object distance (FOD) and 225 mm focus-to-detector distance (FDD), using 40 kV, 300 μA, and 0.5 s exposure for 1400 images, with a frame average of 3 and image skip of 1. This configuration allowed us to decrease the width of the detector to 2,000 pixels, meaning that each X-ray image was 9.15 MB. The Y-axis of the stage was shifted down by 29 mm between scans to allow some overlap between segments (Fig. 3a). For each segment this resulted in a scan time of 47 minutes and 17.8 μm voxel size, with reconstructed volumes around 10 GB each (Fig. 3b).

**Figure 3.**
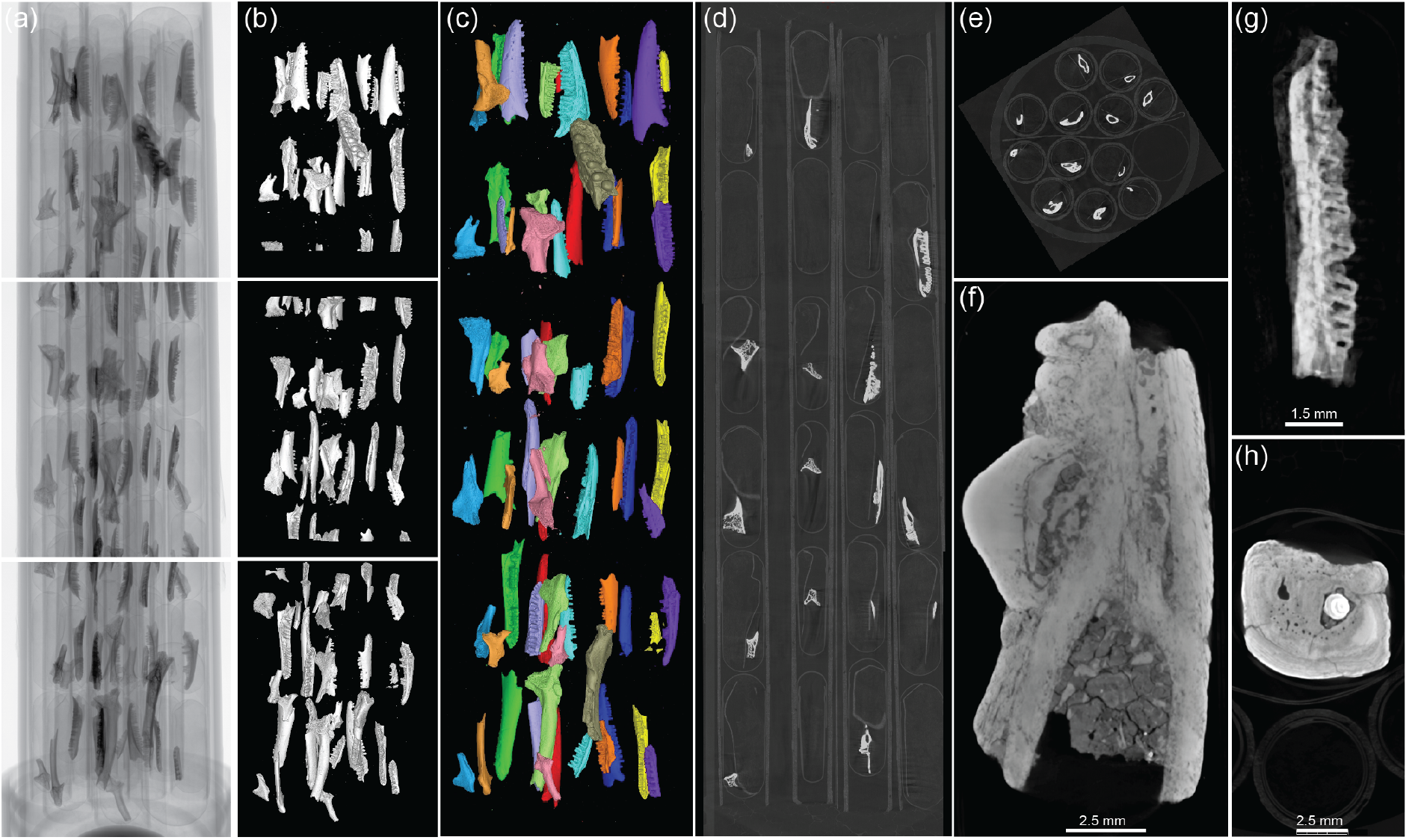
High-throughput μCT results: (a) the first X-ray image from each scanned segment of the 50 ml tube from top (tip) to bottom (cap), containing 72 frog and lizard specimens. (b) 3D renderings of the individual segments in the same positions, at 17.8 μm voxel size, (c) the stitched 3D volume at the same resolution, with fossils colour-coded by straw, (d) reconstructed cross-section of the same tube in side and (e) top views, (f) cross-section of an agamid lizard jaw showing contrast between the matrix, bone, and teeth, (g) cross-section of a skink mandible with a shadow motion artefact, (h) an example of beam hardening caused by a dense surface.

This process is in stark contrast to the effort needed to scan each specimen individually at the highest resolution (10.6 μm over 5,000 projections for a single size 4 capsule), requiring 167 minutes per scan or over eight hours for a fully packed tube, excluding time to swap samples. Automated sample changers are available for some μCT machines (e.g., Bruker SkyScan, Bruker Biosciences Pty Ltd), although these are still limited to under 20 specimens. Reconstructing and rendering each μCT file separately would also require substantial computing time, making the task unfeasible for individual researchers wishing to work on large data sets. Instead, we stitched the three segments together into a volume of equally high resolution (Fig. 3c), allowing access to multiple specimens by opening a single 30 GB file.

The image quality achieved by our method was more than adequate for applications in biological systematics, including geometric morphometrics, functional analyses, and morphological descriptions. Cross-sectional μCT images revealed sharp contrast between the hydroxyapatite of the bone, dentine, and enamel against the clay and crystalline calcium carbonate-rich matrix, while offering precise details of specimen morphology as well as the dividing walls of the capsules, straws and paper (Fig. 3d-f). In few cases artefacts were observed in the reconstruction due to movement (Fig. 3g) or beam hardening (Fig. 3h). These specimens were labelled as ‘REDO’ in the 3D volume and set aside during unpacking to be rescanned, either with additional tissue in the capsule or using a filter, respectively. In earlier scans we observed some capsules in straws that had been slightly crushed by the one above it. These straws were packed using a solid dowel to push down the capsules, hence why we recommend using a cocktail straw instead.

Other examples of material scanned using our high-throughput method are shown in Figure 4. Fossilized frog ilia and varanid osteoderms (bony deposits in the scales of some lizards) yielded excellent results following this protocol (Fig. 4ab), with hundreds of 3D models generated for minimal time, money and effort. We also applied our system to larger mammalian specimens, for example fossilized rodent jaws from the Middle Pleistocene Mt Etna^42^. This material is larger and denser than the herpetological samples, requiring slightly different parameters for μCT scanning. For larger/denser fossils, a 0.1 mm copper plate was secured under the collimator to reduce beam hardening and other artefacts, and X-ray voltage was increased to 50-80 kV. The resulting reconstructed images could be easily segmented using density-based approaches in VGStudioMax, like the region growing tool to separate teeth from bone (Fig. 4c).

**Figure 4.**
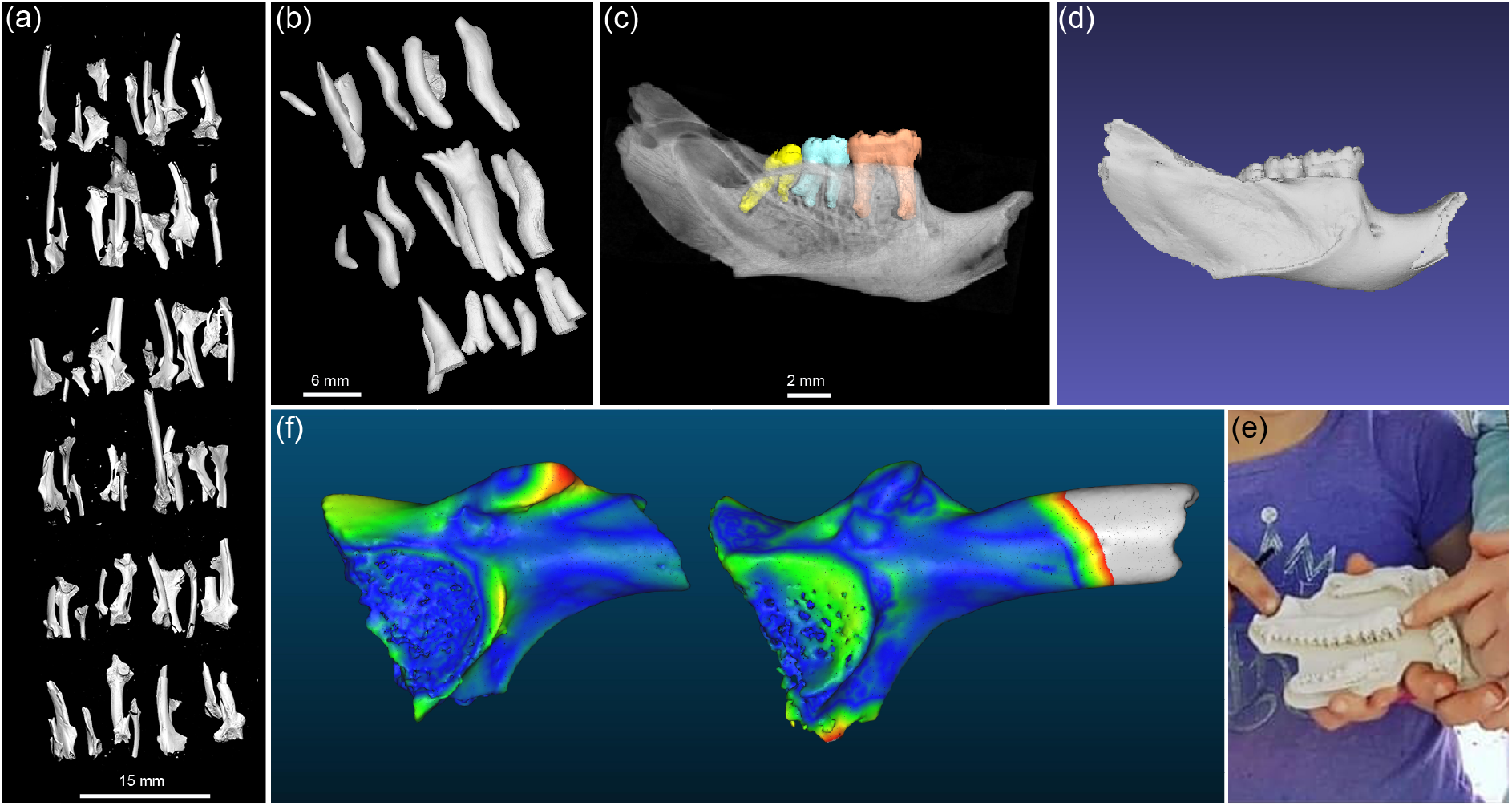
Scanned fossil material and its applications: (a) 3D rendering of 73 fossil frog ilia scanned in a single tube, (b) a subsample of varanid osteoderms in 3D, (c) mandible of the rainforest rodent *Pogonomys* from the Riversleigh World Heritage site with molars colour coded and jaw bone rendered semi-transparent, (d) surface mesh of the same specimen composed of 109,213 vertices (file size 10.5 MB), (e) photograph of children holding a 3D print of enlarged fossil skink jaws at Capricorn Caves Fossil Open Day, (f) point cloud comparison of fossil tree frog (Hylidae) ilia found in Capricorn Caves at 10-20 cm depth (left) and 40-50 cm depth (right), representing a time difference of 2-3,000 years. Warmer colours denote regions with greater shape differences, as estimated in CloudCompare v.2.10.2.

Stewardship of μCT data is still a major challenge for natural history museums, which are now tasked with the curation of rapidly expanding digital datasets associated with the physical collections^43^. There is currently no standard for 3D data storage, meaning that the fate of μCT images and their metadata is often individual or institution-driven. We retain at least two copies of the original X-rays and reconstructions in different locations (one cloud-based, one physical), while working on the prepared volume on a regularly backed-up server. Davies et al.^5^ provide a summary of file types that should be retained for 3D digital morphology analyses, with best practices recommended when storage space is not a problem. Online data repositories are another attractive option for long-term data storage, several of which are free and cater to 3D digital datasets generated for biological research^5^, e.g., Zenodo (zenodo.org), MorphoSource (www.morphosource.org).

For most downstream analyses, μCT data will be converted into polygonal surface models such as STL (sterolithography) files, which capture the geometric shape of a 3D object using triangles or vertices to describe its surface (Fig. 4d). These models, also known as surface meshes, are virtually interactive and can be manipulated, viewed, and measured using the open source software MeshLab (http://meshlab.sourceforge.net/). They are also small in terms of data storage, making them easy to distribute online and providing wider access to museum collections. Finally, surface meshes can be scaled up for displays and educational programs, for example using 3D printing. Following digital labelling and unpacking of our specimens, we extracted STL models of each fossil from its ROI in VGStudioMax, while retaining its original identity.

Many of the fossils included in this project were excavated from cave deposits at Capricorn Caves in Rockhampton, Queensland, Australia. As part of an annual science communication event called Capricorn Caves Fossil Open Day^44^, we 3D printed select specimens to engage the public in local biodiversity. There palaeontologists referred to the enlarged 3D models to assist in the interpretation of microfossils, allowing children to interact with replicas of local animals from thousands of years in the past (Fig. 4e). In addition to public outreach, our future work will focus on analysing the generated μCT data using a growing toolkit of bioinformatic approaches, including machine learning and AI, 3D landmark-based geometric morphometrics, and high-density point cloud comparisons of fossil and living forms (Fig. 4f).

## Conclusions

Current research on museum-based μCT data encompasses an extraordinary array of material and topics, including tetrapod origins^45^, echolocation in bats^46^ and whales^47^, limb reduction in lizards^48,49^, mammalian limb development^50,51^, primate neuroanatomy^52,53^, reproduction in insects^54^ and plants^55^, paleoecology of Precambrian biota^56^, seafloor biomass estimation^57^ and amber inclusions^58^, to name a few. Typically these studies involve few specimens, due to the challenges of capturing high resolution 3D data for large sample sizes.

Here we demonstrate how simple, inexpensive equipment like drinking straws and pharmaceutical capsules can be modified for high-throughput μCT scanning of dozens of fossils in a single container, providing large-scale morphological datasets for comparative analyses. To date we have scanned over 1,000 accessioned museum specimens following this technique, including hundreds of frogs, lizards, snakes and mammals. These data will contribute to ongoing efforts to identify evolutionary responses to climate change in Australia’s fossil record, where reptiles and amphibians constitute half of the terrestrial vertebrate diversity, and where increasing aridity has transformed large parts of the continent. This method can also be adapted for other (non-biological) objects, by creating custom-made holders and stages, and seeking μCT scanners with differing capacities. Some users may wish to scan individual specimens or holotypes at the highest possible resolution, in which case our protocol offers an efficient pathway for determining which specimens should be elevated to type status. At the same time we note that fossils can be difficult to scan, being highly variable in shape, composition, and density. Thus while μCT is a powerful tool for most small specimens, in some cases other non-destructive techniques such as magnetic resonance imaging (MRI), synchrotron, and confocal microscopy may be more appropriate.

## Acknowledgements

This work was supported by an Australian Research Council DECRA (DE180100629) to CAH, with scans performed through the Melbourne TrACEES platform. Danielle Measday, Conservator of Natural Sciences at Museums Victoria, advised on and provided archival papers for testing. We thank Kristen Spring, Rochelle Lawrence, Joanne Wilkinson, and Jonathan Cramb, along with many volunteers at the Queensland Museum, for their assistance in generating the large quantity of specimens made available for this project.

## Author Contributions

C.A.H. and S.H. conceived the project; S.H. selected specimens for μCT scanning; C.A.H., R.A., and J.R.B. designed the methodology; J.R.B. operated the μCT machine; C.A.H. and R.A. collected and analysed the data and led manuscript writing. All authors contributed to manuscript drafts and gave final approval for publication.

## Data Availability

All X-ray and μCT data shown in Fig. 3a-e are available in the MorphoSource repository, [link provided upon acceptance].

